# Real-time object locator for cryo-EM data collection --- You only navigate EM once ---

**DOI:** 10.1101/2021.04.07.438905

**Authors:** Koji Yonekura, Saori Maki-Yonekura, Hisashi Naitow, Tasuku Hamaguchi, Kiyofumi Takaba

## Abstract

In cryo-electron microscopy (cryo-EM) data collection, locating a target object is the most error-prone. Here, we present a machine learning-based approach with a real-time object locator named yoneoLocr using YOLO, a well-known object detection system. Implementation showed its effectiveness in rapidly and precisely locating carbon holes in single particle cryo-EM and for locating crystals and evaluating electron diffraction (ED) patterns in automated cryo-electron crystallography (cryo-EX) data collection.

---

Cryo-EM work can extend over hours or days during which thousands of image stacks are collected for high-resolution single particle reconstructions. Automated data collection with minimal human-supervision would be advantageous. A number of computer programs have been developed to fulfill this (e.g. refs. 1 - 3), and are widely used in laboratories and shared facilities. Automation is also becoming increasingly important in electron 3D crystallography / 3DED / microED, where acquisition of rotational diffraction data from many small crystals is often needed (e.g. refs. 4, 5).

The operator’s tasks in single particle analysis (SPA) data collection typically comprise: alignment of the microscope; selection of grid squares and carbon holes containing sample molecules from low magnification images; and start data acquisition. Data acquisition requires expediting search, focus and record modes. In search mode, a target carbon hole with a diameter of 1 - 2 μm and filled with amorphous ice is positioned at intermediate magnification. This maneuver is the most error-prone due to discrepancies in stage positions between low and intermediate magnifications and a poor positional reproducibility or mechanical instability of the specimen stage. The stage rarely goes to an exact pre-registered position within an allowable error range, typically, a few hundred nano-meters. Precise adjustment of the stage position in search mode is critical and yet often frustrating. Stage alignment to a hole is done by cross-correlating a test image at 1,000 to 10,000× magnification and a reference, an ideal hole image at the same magnification or a corresponding image at lower magnification, prepared beforehand. However, this step can be problematic, particularly when poor signals in thick ice, incorrect or broad correlation peaks of partially recorded multiple holes, and contamination.

Modern schemes take 9 – 25 or more image stacks from holes clustered around a central hole and / or even more stacks for one hole through changing deflector coils once the stage is moved to a new position ^6^. This approach is indeed very effective to speed up data collection, but, when the alignment fails, it can be disastrous, yielding much useless data, wasting time and data storage space. To avoid this, EM operators can try higher defocus values, longer exposure times, larger frame binning, and, if applicable, filtration of inelastically scattered electrons ^7^ for gain in image visibility. However, these interventions require operator input, is time-consuming, and can be ineffective in difficult cases. Also, during unsupervised data acquisition multiple trials for positioning one hole may be repeated, resulting in possible radiation damage.

Instead of relying on cross correlations, we have developed a new algorithm incorporating machine-learning and the well-known real-time object detection system YOLO (You Only Look Once) ^8^. Machine-learning has already been introduced in the cryo-EM field to solve various problems in data analyses such as particle picking (e.g. refs. 9 - 11), denoising (e.g. ref. 12), analysis of structure variations (e.g. ref. 13) and so on. However, it has not, or has hardly, been used in the control and supervision of data collection to our knowledge, while YOLO was originally designed for real-time object detection. Our software named yoneoLocr (You Only Navigate EM Once to LOCate objects in Real time) is based on the latest release of YOLO version 5 (YOLOv5; https://github.com/ultralytics/yolov5), and can detect objects in less than 0.1s using trained weights.

Implementation required using cryo-EM images containing carbon holes recorded at a nominal magnification of 8,000× on a Gatan K3 direct electron detection (DED) camera with a JEOL CRYO ARM 300 electron microscope ^14, 15^. This magnification is generally used for stage alignment in search mode of SPA data collection, but here the in-column type energy slit is retracted due to a large cut in the view. We then enclosed the carbon holes in these images with a box and annotated them as “hole”. The annotated images were trained with a network model of YOLOv5s on 4× GPU cards installed on a Linux workstation. Hole detection was excellent. Unsurprisingly, YOLOv5 out-performed YOLOv3, and the simplest model YOLOv5s showed an excellent performance for hole detection.

We made a python program named yoneoLocrWatch.py (Fig. 1) to monitor updates in a predefined directory of a K3 control Windows PC equipped with two GPU cards. One GPU card may be enough as the computation and memory loads are low during running of the object detection routine and at other times the program is idle. Once a text file in the directory is renewed, the program reads an image indicated in the text file and provides the best directions for stage shifts based on confidence level, hole size, and distance from the center of the image. Hole detection typically takes ∼0.06s. We made a SerialEM script, AlignyoneoHole, for incorporating this hole detection step into automated data acquisition (Fig. 1). The script takes a new image, updates the directory, reads a log, and shifts the stage if needed. This approach never failed in stage alignment, even for difficult samples with large contamination and less visibility of thick ice or carbon film over holes, as long as holes were recognizable by the human eye (Figs. 2a - d), in notable contrast to the cross correlation method (Figs. 2a and b). In one dataset collected using yoneoLocr from a holey carbon grid covered with a thin carbon layer, ∼ 91% of images were correctly aligned within two trials and the remaining ∼ 9% were aligned by the third trial. The success rate within second trials reached ∼ 94% for typical samples on standard holey carbon grids or gold-sputtered holey carbon grids ^14^. Obviously there are additional advantages - ∼5 - 50-fold less exposure time (decreased from 0.5 – 2 s to 0.03 - 0.2 s for taking one search image and requiring less repeats till positional error is small enough), therefore less radiation damage, and no or less defocusing, therefore minimal objective lens changes and more stable data collection. A typical job for hole detection is shown in Supplementary Video 1.

**Figure 1.**
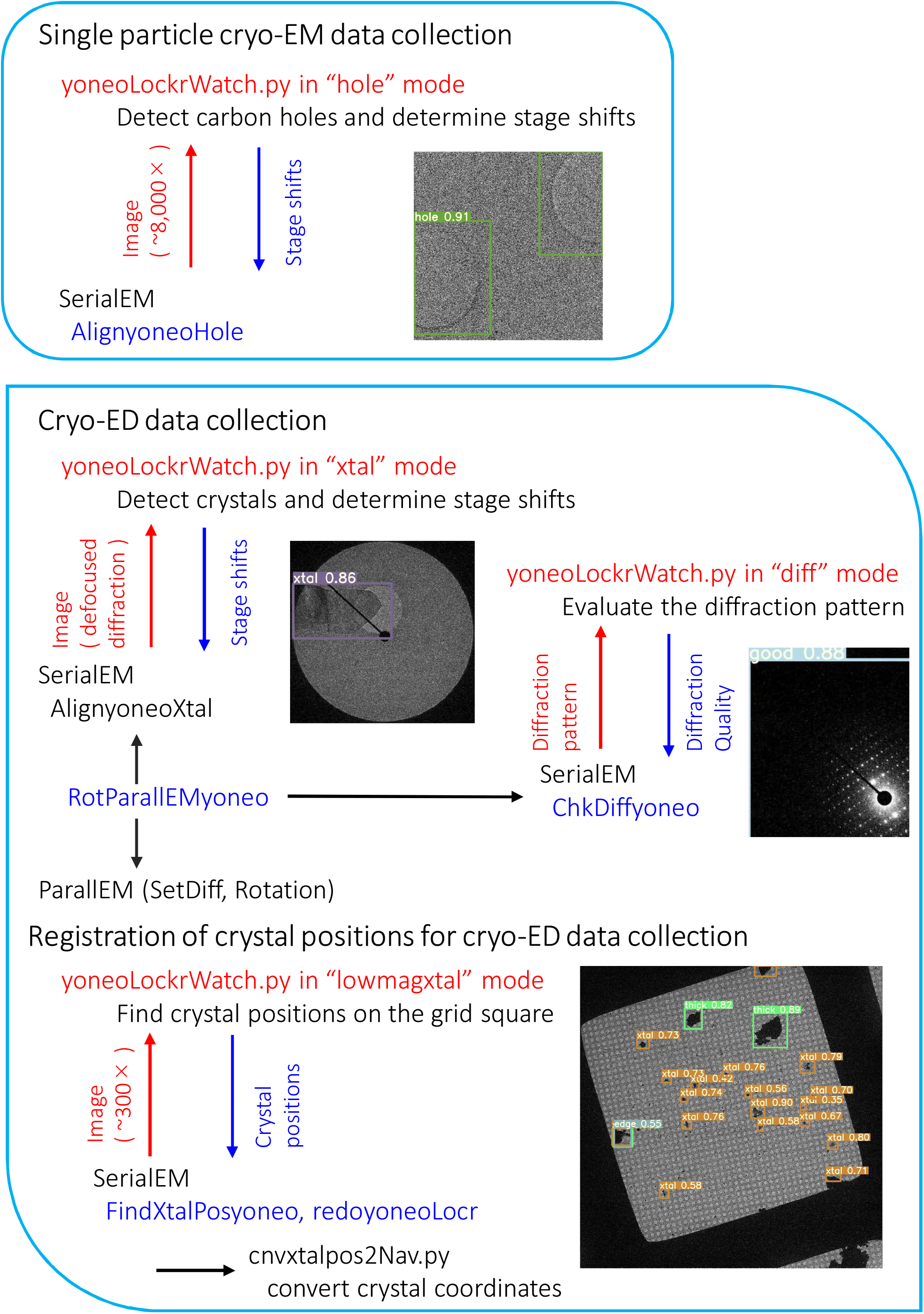
**Schematic diagram for positioning the target and checking the quality of diffraction patterns by yoneoLocrWatch.py, SerialEM scripts and associated programs.**

**Figure 2.**
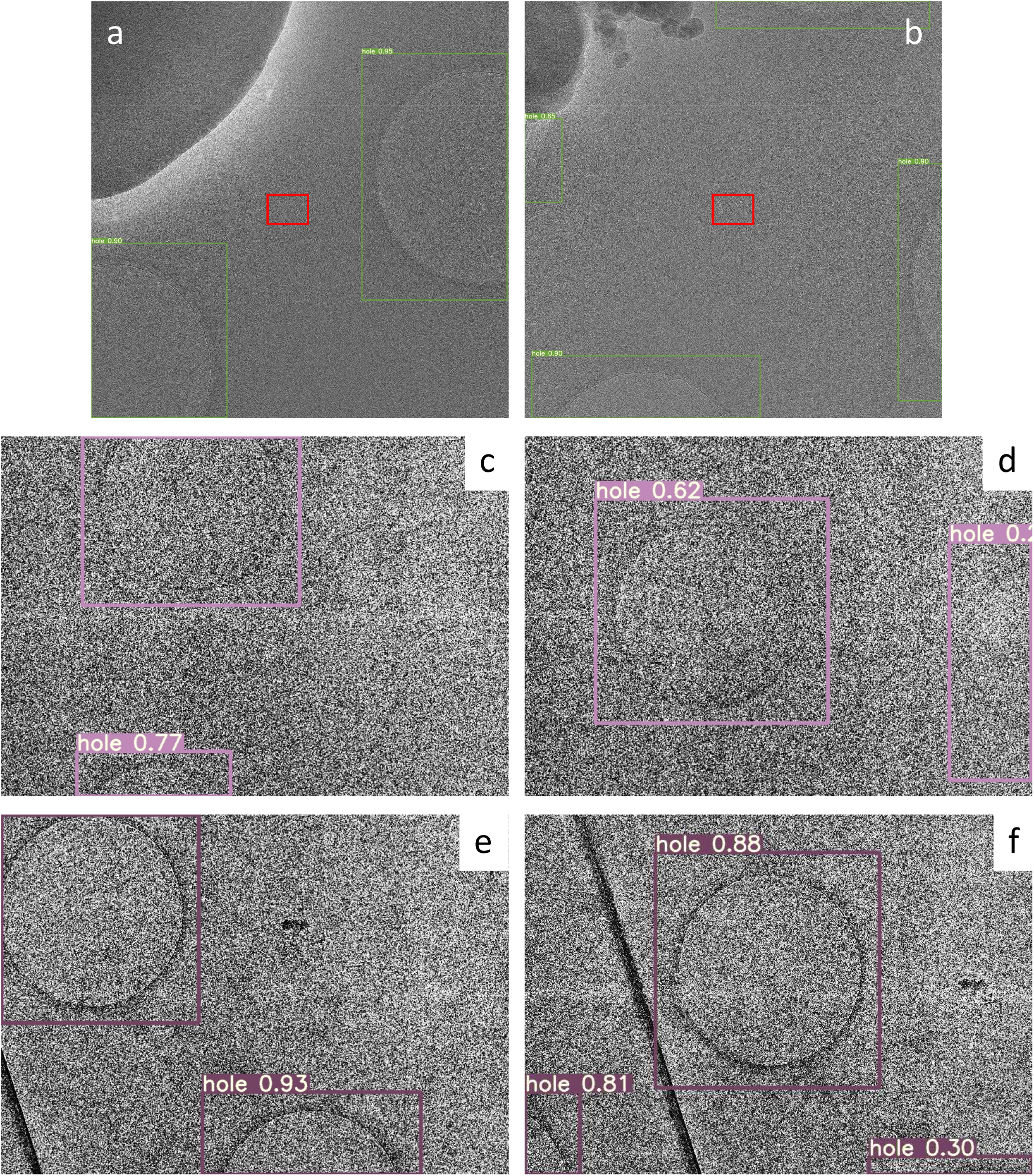
Detection of carbon holes. **a**., **b**. Failures of stage positioning done by cross-correlation with a reference image using SerialEM due to excessive contamination in dark blobs and to partially recorded multiple holes. Center small rectangles in red indicate a rough position and dimension of acquired movie stacks on the K3 camera at a nominal magnification of 100,000×. Thus, each of these alignments produced 25 useless image stacks, as image shifts were applied over 5 × 5 neighboring holes for data collection at that time. YOLOv5 with a weight trained in this study detected all holes (boxes) correctly. **c**., **d**. Low-contrast images of holes covered with a thin carbon layer and with thick ice. The trained weight still succeeded in hole detection (boxes). **e**. The first image taken in Search mode. **f**. The second image after stage adjustment by yoneoLocrWatch.py running in hole mode and AlignyoneoHole.

There have been several reports and programs for automated collection of rotational ED data (e.g. refs. 16, 17). Our group also developed and reported a scheme that combines SerialEM for positioning of sample crystals and ParallEM for controlling data acquisition ^18^. This protocol needs manual positioning of crystals and registration of the coordinates in defocused diffraction (search) mode before unsupervised data acquisition is started in focused diffraction (data-taking) mode. Thus the same transposition problem seen in SPA data collection arises. Since the appearances of crystals differ from crystal to crystal, it is more challenging than for hole detection. The original protocol does not include re-positioning to target crystals during data collection and is more severely affected by positional reproducibility.

YOLO was used again for training with various crystal images of proteins ^19^, polypeptides, organic molecules ^5^, and semiconductor materials ^4^ recorded in defocused diffraction mode on different cameras, two scintillator-coupled CMOS cameras, Gatan OneView ^19^ and TVIPS XF416 ^5, 15^, and one DED camera Direct Electron DE64 ^4^. As samples often contain ice crystals of characteristic hexagonal shape and yielding strong hexagonal diffraction patterns, crystal images were annotated as “xtal” or “ice” according to appearance. We also trained YOLOv5 with ED patterns of crystals acquired in data-taking mode. Diffraction patterns were visually assigned as “good”, “soso”, “bad”, “no” and “ice” depending on the quality of the patterns within a single box covering most of the whole image. The worst level “no” had patterns with no diffraction spots. The most accurate model (YOLOv5x) was adopted for both crystal images and diffraction patterns.

Runs, incorporating the trained weights, were implemented in DE64 and XF416 camera control PCs equipped with GPU cards. Two sessions of yoneoLocrWatch.py were launched simultaneously in two different modes, “xtal” for locating a crystal and “diff” for evaluating the diffraction pattern (Fig. 3). Two SerialEM scripts, AlignyoneoXtal and ChkDiffyoneo, were made for incorporation of these operations into script RotParallEMyoneo that calls up ParallEM for sequential data collection of rotational diffraction patterns in a queue list in the Navigator window of SerialEM (Fig. 1). SetDiff in ParallEM switches between defocused (search) and focused (data-taking) modes by changing only the intermediate lens 1 (IL1) value. During data collection, it never goes back to imaging nor low magnification modes. Thus, possible hysteresis from lens changes is minimal and the beam setting is stable even over operations taking several days, which could include overnight breaks with the beam off and multiple flashing cycles for refreshing the cold-field emission gun in the CRYO ARM 300 microscope ^14, 15, 18^. SetDiff also controls insertion and retraction of the selected aperture and energy slit ^15, 18^.

**Figure 3.**
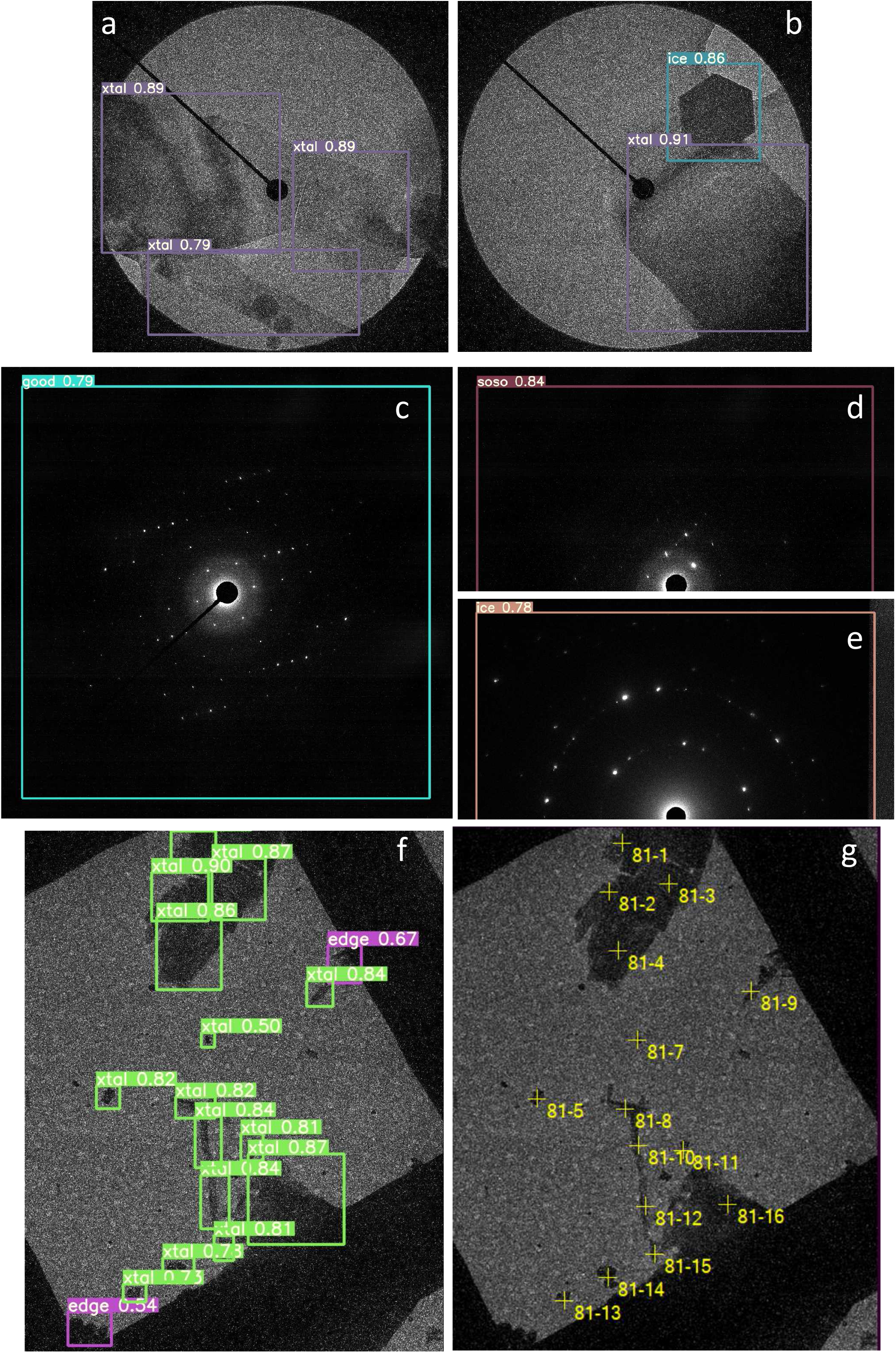
Detection of crystals and check of diffraction patterns. **a**., **b**. Detection of crystals in defocused diffraction mode by yoneoLocrWatch.py in xtal mode. A hexagonal ice crystal was identified in b. **c**., **d**., **e**. Evaluation of diffraction patterns by the same python program in diff mode. A diffraction pattern from an ice crystal was identified in e. **f**. Detection of crystals taken in a low-magnification (300×) image in lowmagxtal mode. Crystals at the square edge were also identified. **g**. Registration of crystal positions to a queue list in SerialEM by FindXtalPosyoneo and cnvxtalpos2Nav.py. The crystals at the edge were automatically excluded in this case.

Thus, identifying suitable crystals and positioning one to center view are possible in defocused diffraction mode, as is done for carbon holes in the SPA data collection scheme (Figs. 3a and b). This step mostly performed well, but, different from regular hole patterns, moving to the best crystal sometimes failed when the crystals are, for example, crowded or too thick. Thus, a prior quality check of untilted still diffraction patterns would be helpful, as acquisition of one rotational data set takes 2 - 3 minutes and rotational frames fill large disk space. This can be done in data-taking mode with a short exposure of ∼0.1s (Fig. 3c; Supplementary Fig. 1). It works particularly well for protein and organic semiconductor crystals, which yield many diffraction spots. Data taking with other crystals succeed at the low assessment levels “soso” or even “bad” (Supplementary Fig. 1). The worst level “no” assigned for no diffraction spots was always easily determined for all samples tested in this study (Supplementary Fig. 1c). Once the diffraction pattern is judged to be above a given level, RotParallEMyoneo calls up Rotations in ParallEM to start rotational data acquisition (Fig. 1).

Finally, we annotated crystals in single grid squares in low-magnification images (250 - 400×) and trained with YOLOv5s again. We categorized crystals as “xtal” for candidates for data collection, “thick” for dense samples, and “edge” for those at the periphery of the grid square roughly normal to the rotation axis. Data collection from crystals at the edge may be blocked by the grid bar at high tilt angles. Thus, the approximate positions of crystals can be obtained by yoneoLocrWatch.py running in the new mode “lowmagxtal”, and SerialEM script, FindXtalPosyoneo, which calls up a python script, cnvxtalpos2Nav.py, to convert crystal coordinates, and adds the positions to a queue list (Figs. 1, 3f and g). This step can be retried with a new confidence threshold level for crystal detection by redoyoneLocr without retaking images. Users may also edit the queue list. Now, registration of suitable crystals and unsupervised rotational data collection are almost fully automated by combining the three implementations in yoneoLocr and associated programs above. Automatic positioning to crystals can be implemented in several published programs, which set certain threshold pixel values to define crystal areas ^16, 20^, but they cannot discriminate ice crystals, nor avoid thicker ones, and do not check for quality.

In summary, we have developed a real-time object locator yoneoLocr for cryo-EM data collection based on machine-learning, which obviates development of special algorithms for each target. Application of the software with SerialEM scripts is effective in efficient unsupervised data collection for SPA and electron 3D crystallography. Object detection is very fast, and can be done with small CPU and GPU loads and minimal memory. There is no delay in running SerialEM and ParallEM, usage is simple, and the routine can be called from other software. Locating carbon holes is extremely precise, with short exposure time, and is even effective in hitherto difficult cases – greatly benefiting SPA data collection. Implementation for automated cryo-EX data collection also performs well, but may need further training with more ED data. The program is also able to re-tune the weight with new data during data collection, which is a function implemented in YOLOv5. We have yet to thoroughly test this feature. The software yoneoLocr including SerialEM scripts and associated programs can be obtained from a GitHub site (https://github.com/YonekuraLab/yoneoLocr).

## Acknowledgments

We thank D. B. McIntosh for help in improving the manuscript. This work was partly supported by JST-Mirai Program Grant Number JPMJMI20G5 (to K. Y.), JST CREST Grant Number JPMJCR18J2, Japan (to K.Y., S. M.-Y., and K. T.), and the Cyclic Innovation for Clinical Empowerment (CiCLE) from the Japan Agency for Medical Research and Development, AMED (to K.Y.).

## Methods

YOLOv5 (https://github.com/ultralytics/yolov5) was installed under python 3.8 environment created by Anaconda in a linux work station equipped with 4× NVIDIA GeForce Titan X GPU cards. This version of YOLO uses PyTorch for deep learning together with CUDA, cuDNN, and so on. A real-time object locator yoneoLocrWatch.py developed in this study was set up under the same environment created by Miniconda or Anaconda on camera control PCs operated by Microsoft Windows Server 12R for K3, and Windows 10 for DE64 and XF416 cameras. The program and associated scripts were placed in C:¥ProgramDat¥ in the PCs. The K3 control PC is equipped with two GPU cards, Quadro K2200 and P6000, the DE64 PC with two Quadro RTX6000s and the XF416 PC with a Quadro K420 and a P4000. ImageMagick was used for pretreatment of images in yoneoLocrWatch.py. Details of the installation and command line options are presented in Supplementary method.

Cryo-EM images and ED patterns in jpeg format were annotated with GUI software labelImg.py (https://github.com/tzutalin/labelImg), and the format of coordinates and class names for annotated objects were converted to the YOLO format using convert2Yolo.py (https://github.com/ssaru/convert2Yolo). All training was done on the linux workstation using train.py included in YOLOv5. Detection test was performed with detect.py.

For hole images, several variations of original images were created by applying binning of images and/or histogram equalization using convert of ImageMagick. Training for the images took ∼1.9 h using a network model YOLOv5s with an image size of 800 × 800. The most accurate model (YOLOv5x) was used for training of crystal images taken in search mode and diffraction patterns in data-taking mode. It took ∼ 4 h with an image size of 800 × 800 and ∼ 20 h with an image size of 1024 × 1024. Training of crystals taken in low magnification images was done using the YOLOv5s model.

For control and unsupervised data acquisition, yoneoLocrWatch.py keeps watching updates in a predefined directory, WatchHole¥ in hole mode, WatchXtal¥ in xtal mode, WatchDiff¥ in diff mode and WatchLowmagXtal in lowmagxtal mode. Once a text file in these directories is renewed, the program reads an image indicated by the text file. Running in hole mode can enhance the image contrast by applying binning and histogram equalization. Then, it detects holes and gives the best answer for stage shifts in a log file based on the confidence level, the hole size, and distance from the center of the image. Running in other modes also gives the results of detection or evaluation in a log file.

SerialEM scripts AlignyoneoHole, AlignyoneoXtal, ChkDiffyoneo, and FindXtalyoneo take images, put a text file containing the location and name of a newly recorded image in the directories yoneoLocrWatch.py is watching, and obtain alignment parameters or quality check from a log file of yoneoLocrWatch.py. Then, AlignyoneoHole and, AlignyoneoXtal moves the stage if needed. As the pixel size cannot be defined in diffraction mode, we spread the beam to the detector edge and supply the beam size (∼ 5 or ∼ 7 μm) to AlignyoneoXtal as a scale for stage shifts. ChkDiffyoneo determines whether the corresponding crystal is worth rotational data collection or not. FindXtalPosyoneo finds crystals in low magnification images and adds the crystal position in a queue list in the Navigation window of SerialEM. This crystal detection can be retried with a new threshold of the confidence level by a SerialEM script redoyoneoLocr without taking the same images again. The retry needs to stop yoneoLocrWatch.py running in lowmagxtal mode before starting the job. Please also refer to Fig. 1 for the workflows.

## Supplementary Information

## Supplementary methods

### Installation

1. Download yoneoLocr-main.zip from https://github.com/YonekuraLab/yoneoLocr.
2. Extract the zip file and put the whole directory as yoneoLocr in C:¥ProgramData¥ of a camera control Windows PC.
3. Set the property of batch files to “full control” from the Security tab if needed.
4. Install CUDA toolkit 10.1 and cuDNN 10.1 for a K3 control PC if the operating system of the PC is Windows Server 12R. CUDA 10.1 is the newest version supporting Windows Server 12R. Newer versions of CUDA and cuDNN are available for Windows 10.
5. Install Microsoft Build Tools for Visual Studio (vs_buildtools) if needed.
6. Install ImageMagick.
7. Launch Anaconda Prompt. Make and activate an Anaconda environment as, conda create -n yolov5-4.0 python=3.8 conda activate yolov5-4.0
8. Go to the yoneoLocr-yolov5 directory and install python libraries as, conda install -c pytorch torchvision cudatoolkit=10.1 pip install -r requirements.txt All required libraries are written in requirements.txt.
9. Make shortcuts of yoneoHole.bat, yoneoXtal.bat, yoneoDiff.bat, and yoneoLowmagXtal.bat on the desktop.
10. Launch yoneoLocrWatch.py from the shortcuts.

### Command line options

--object hole / xtal / diff / lowmagxtal

Select running mode.

--conf-sel 0.4

A confidence threshold for object selection in hole and lowmagxtal modes.

Default 0.4.

--delout yes / no

Delete output file showing objects enclosed with boxes. Default: no.

--ice yes / no

Include ice crystals for positioning in xtal mode. Default: no.

Other options in the original script detect.py in YOLOv5 are also available.

**Supplementary Fig. 1.**
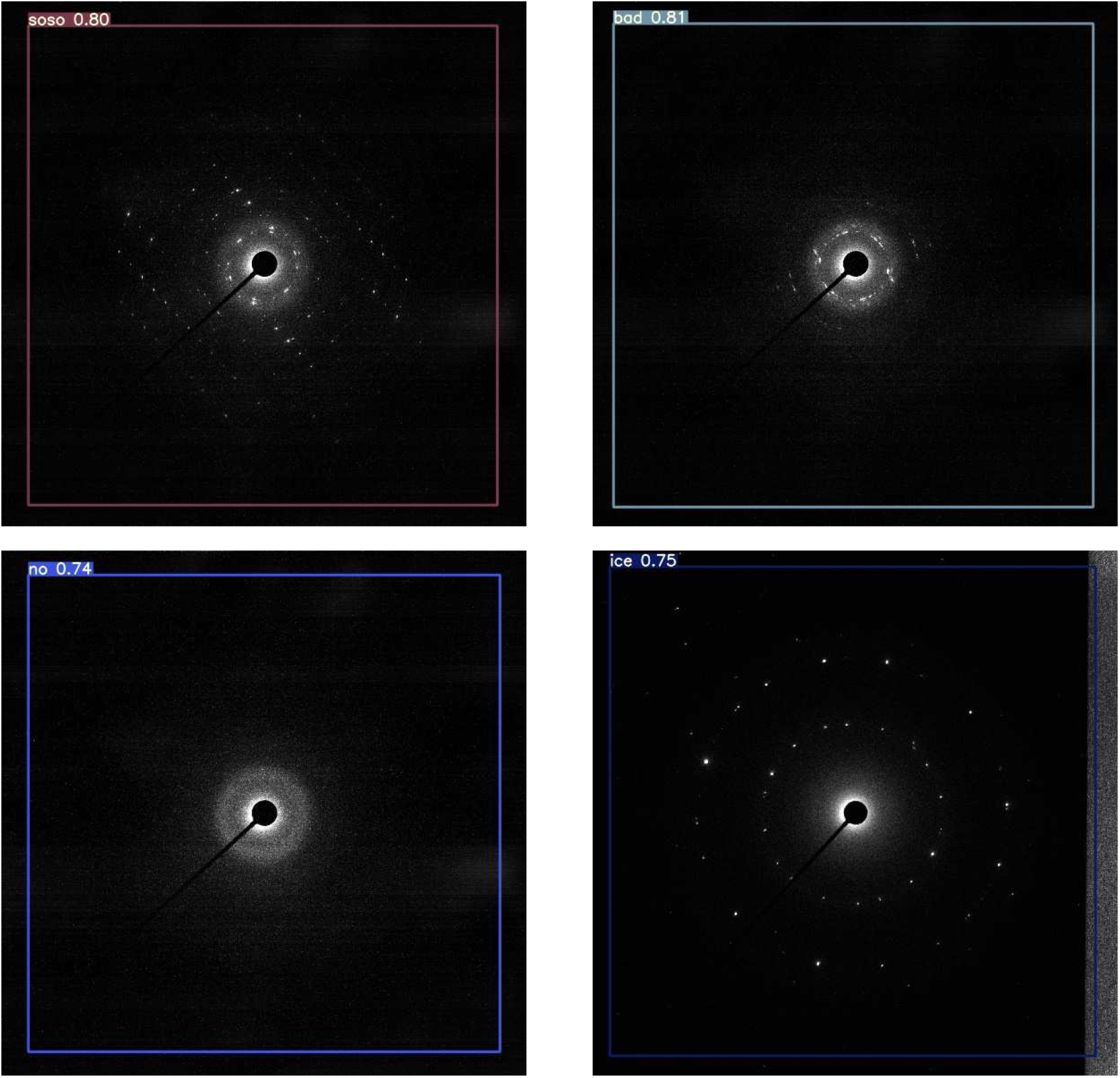
Typical evaluation results of diffraction patterns.

